# *Brca1* heterozygosity leads to hepatic steatosis in male and female mice despite sexually dimorphic effects on systemic metabolism

**DOI:** 10.64898/2026.02.25.708005

**Authors:** Sailesh Palikhe, Linlan Qiao, Caleb Kutz, Xiaobo Liang, Poojaben Dhorajiya, Devin C. Koestler, Priya Bhardwaj, Colin S. McCoin, Orion Peterson, Hubert M. Tse, John P. Thyfault, Kristy A. Brown

## Abstract

Carrying a germline mutation in *BRCA1* is associated with an increased risk of several cancers, including breast and ovarian. Our recent work has demonstrated that obesity is associated with elevated levels of DNA damage in breast glands in this high-risk population. *BRCA1* is a canonical tumor suppressor gene primarily recognized for its role in DNA damage repair, yet emerging evidence suggests broader functions in metabolic regulation. To determine whether heterozygous loss of *Brca1*, as seen in individuals who carry a germline mutation, modifies susceptibility to diet-induced metabolic dysfunction in a sex-dependent manner, we subjected wild-type (WT) and *Brca1*^+/−^ mice of both sexes to a high-fat diet (HFD) and performed longitudinal metabolic phenotyping. Female *Brca1*^+/−^ mice exhibited pronounced obesity, increased adiposity, hyperinsulinemia, and impaired glucose tolerance. In contrast, male *Brca1*^+/−^ mice showed modest resistance to HFD-induced weight gain and displayed improved glucose tolerance compared to WT controls. Notably, *Brca1* heterozygosity led to more severe hepatic steatosis with HFD, indicating a shared susceptibility to liver lipid accumulation despite divergent systemic outcomes. In females, steatosis was associated with reduced mitochondrial respiratory complex IV activity and transcriptional remodeling that favored lipid storage. Treatment with the dual GLP⍰1/GIP receptor agonist tirzepatide ameliorated systemic metabolic dysfunction and hepatic steatosis in HFD–fed female *Brca1*^+/−^ mice. These findings identify *Brca1* heterozygosity as a modifier of metabolic disease risk, expanding *BRCA1* biology beyond tumor suppression.

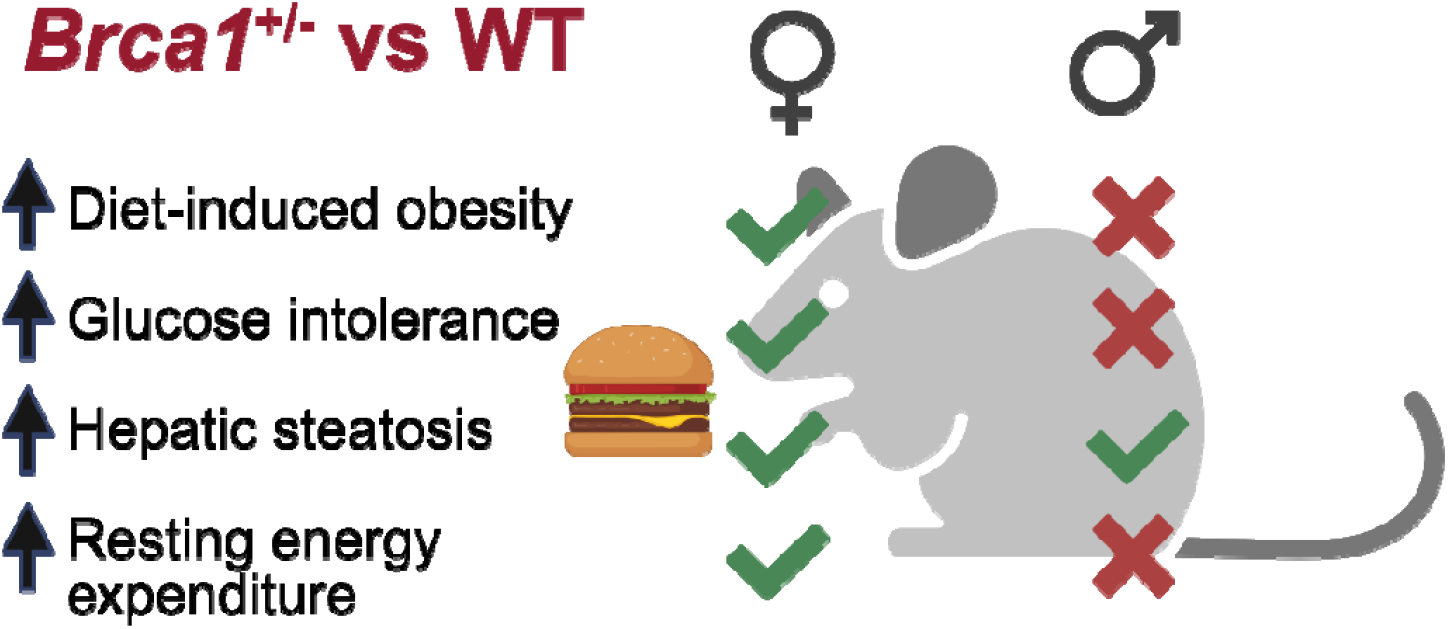

## Introduction

The breast cancer susceptibility gene *BRCA1* encodes a tumor suppressor protein essential for DNA damage repair and maintenance of genomic stability^1,2^. Pathogenic germline mutations in *BRCA1* are typically inherited in a heterozygous state and confer markedly increased breast cancer risk^3,4^.

Metabolic dysregulation has been increasingly linked to breast cancer risk and progression, including in *BRCA1* mutation carriers^5-10^. Clinical studies have reported a higher prevalence of diabetes among *BRCA1 or BRCA2* (*BRCA1/2*) mutation carriers following breast cancer diagnosis^11^, and carriers of *BRCA1/2* loss-of-function variants exhibit elevated fasting glucose and insulin levels compared with carriers of variants of unknown significance^12^. Consistent with these observations, we previously demonstrated that obesity exacerbates DNA damage and mammary tumor penetrance in female *Brca1* heterozygous knockout mice (*Brca1*^+/−^) fed a high-fat diet (HFD), a model that recapitulates key features of human pathogenic *BRCA1* mutation carriers^10^. Together, these findings raise the possibility that *BRCA1* heterozygosity alters systemic metabolic homeostasis, thereby contributing to disease susceptibility.

Here, we tested the hypothesis that *Brca1* heterozygosity increases vulnerability to diet-induced metabolic dysfunction. Wild-type (WT) and *Brca1*^+/−^ mice of both sexes were maintained on a low-fat diet (LFD) or HFD for 22 weeks, and metabolic parameters were assessed longitudinally. We observed marked sexually dimorphic effects across most metabolic endpoints in response to HFD. Female *Brca1*^+/−^ mice developed exacerbated diet-induced obesity, increased adiposity, and impaired glucose tolerance compared with WT controls. In contrast, male *Brca1*^+/−^ mice were partially protected from HFD-induced weight gain and exhibited improved glucose tolerance relative to WT males.

Despite these opposing systemic phenotypes, hepatic steatosis was more severe in both female and male *Brca1*^+/−^ mice, revealing a sex-independent role of *Brca1* in maintaining liver metabolic homeostasis. This finding is particularly notable given the relative resistance of female mice to HFD-induced hepatic steatosis^13^, underscoring a previously unrecognized role for Brca1 in protecting the liver from lipid accumulation. Finally, we show that tirzepatide, a dual glucagon-like peptide-1 (GLP-1) and glucose-dependent insulinotropic polypeptide (GIP) receptor agonist, effectively ameliorates HFD-induced metabolic dysfunction in female *Brca1*^+/−^ mice, including hepatic steatosis.

Collectively, these findings extend the biological relevance of *BRCA1* beyond cancer susceptibility to vulnerability to metabolic disease, with potential implications for metabolic screening and preventive strategies in *BRCA1* mutation carriers.

## RESULTS

### 1. *Brca1* heterozygosity promotes high-fat diet-induced obesity and glucose intolerance in female mice but protects male mice

To investigate the impact of *Brca1* heterozygosity in females, WT and *Brca1*^+/−^ mice were fed a LFD or HFD from 4 weeks of age for 22 weeks (**Figure 1A**). Baseline body weight was comparable between genotypes. However, under HFD conditions, female *Brca1*^+/−^ mice gained significantly more weight than WT females (**Figure 1B**; WT (22 weeks): 35.93g ± 2.53 vs. *Brca1*^+/−^ (22 weeks): 42.83g ± 1.68, *p = 0*.*02)*, while LFD-fed females showed no difference. Energy intake was not significantly different between *Brca1*^+/−^ and WT mice (**Figure 1C**).

**Figure 1.**
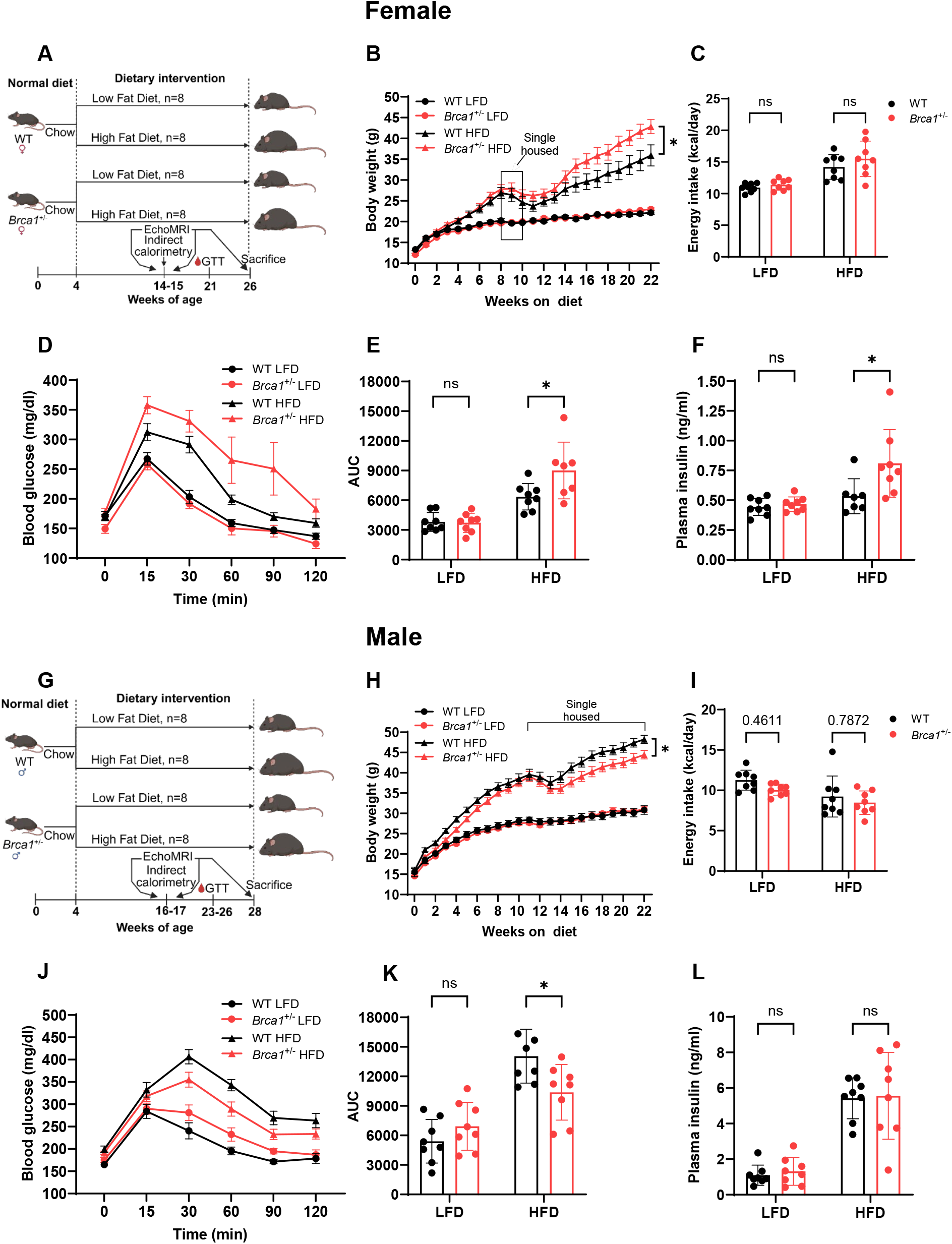
Sex-specific effects of diet and *Brca1* heterozygosity on body weight and glucose tolerance. (A) Experimental design for female mice. (B) Body weight of female WT and *Brca1*^+/−^ mice fed LFD or HFD for 22 weeks (n = 8/group). (C) Energy intake measured during 1-week single housing at 10 weeks on diet (n = 8/group). (D) Glucose tolerance test (GTT) and (E) area under the curve (AUC) in female mice after 17 weeks on diet (n = 8/group). (F) Fasting plasma insulin levels in female mice after 22 weeks on diet (2-hour fast, n = 8/group). (G) Experimental design for male mice. (H) Body weight of male WT and *Brca1*^+/−^ mice fed LFD or HFD for 22 weeks (n = 8/group). (I) Energy intake during 1-week single housing at 10 weeks on diet (n = 8/group). (J) GTT and (K) AUC in male mice after 19–22 weeks on diet (n = 8/group). (L) Fasting plasma insulin levels in male mice after 24 weeks on diet (2-hour fast, n = 8/group). Data are represented as mean ± SEM. \**p* < 0.05, ns, not significant, by two-way ANOVA with Tukey’s multiple comparisons test (B, C, E, F, H, I, K, and L).

HFD-fed female *Brca1*^+/−^ mice also exhibited impaired glucose tolerance (**Figure 1D**), with higher area under the curve (AUC) (**Figures 1E**; WT: 6354.88 ± 471.19 vs. *Brca1*^+/−^ : 9008.86 ± 1084.22, *p* = *0*.*02*), and elevated fasting plasma insulin (**Figure 1F**; WT: 0.53ng/ml ± 0.06 vs. *Brca1*^+/−^: 0.81ng/ml ± 0.10, *p* = *0*.*02*). Glucose-stimulated insulin secretion (GSIS) assays in isolated islets showed comparable insulin secretion between genotypes (**Figure S1A**), indicating that elevated fasting insulin *in vivo* was not due to intrinsic β-cell hyperactivity.

We next examined the effect of diet in male mice using the same experimental design, with male mice fed for 24 weeks (**Figure 1G**). In contrast to females, HFD-fed male *Brca1*^+/−^ mice displayed significantly lower body weight compared with HFD-fed WT male mice after 22 weeks on diet (**Figure 1H**; WT: 48.25g ± 1.01 vs. *Brca1*^+/−^: 44.46g ± 1.05, mean ± SEM, *p* = *0*.*046*), with no changes in energy intake (**Figure 1I**). Moreover, HFD-fed male *Brca1*^+/−^ mice exhibited improved glucose tolerance relative to HFD-fed male WT mice (**Figure 1J**), with a significantly lower AUC (**Figures 1K**; WT: 14053.50 ± 966.79 vs. *Brca1*^+/−^: 10379.75 ± 997.67, mean ± SEM, *p* = *0*.*04*). However, fasting plasma insulin levels and GSIS responses were not different between groups (**Figures 1L and S1B**).

### 2. *Brca1* heterozygosity alters lean mass, adiposity and adipose tissue inflammation in female and male mice fed a high-fat diet

Because HFD-fed female *Brca1*^+/−^ mice exhibited greater body weight, body composition was assessed at 22 weeks using EchoMRI. Fat mass was significantly higher in HFD-fed female *Brca1*^+/−^ mice compared with WT controls (**Figure 2A**; WT: 15.21g ± 2.15 vs. *Brca1*^+/−^ : 20.82g ± 1.21, *p = 0*.*02*), while lean mass was significantly increased (**Figure 2B**; WT: 19.06g ± 0.41 vs. *Brca1*^+/−^ : 20.88g ± 0.42, *p* < *0*.*01*). Early time points (8-10 weeks) showed no differences in fat mass, but lean mass was higher at these time points (**Figures S2A and S2B**).

**Figure 2.**
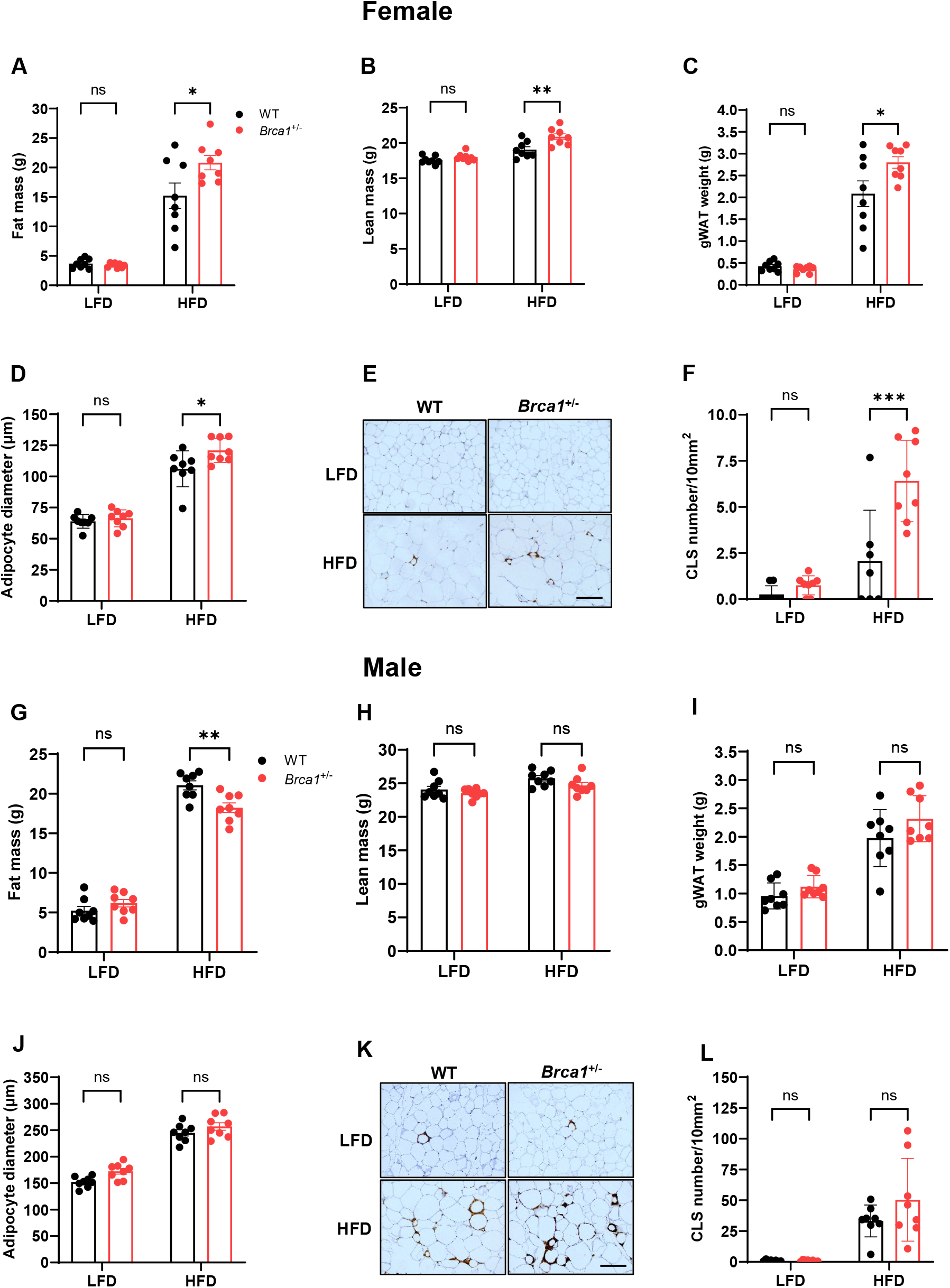
Sex-specific effects of diet and *Brca1* heterozygosity on adiposity and adipose tissue inflammation. (A) Fat mass and (B) lean mass of female mice at 22 weeks, measured by EchoMRI (n = 8/group). (B) Gonadal white adipose tissue (gWAT) weight in female mice after 22 weeks (n = 8/group). (C) Adipocyte diameter in gWAT of female mice quantified from H&E-stained sections (n = 8/group). (D) Representative F4/80 immunostaining in female gWAT (10x, scale bar = 150□μm) (E) Crown-like structures (CLS) quantified by F4/80 staining of female gWAT (n = 8/group; 3 fields per section). (F) Fat mass and (H) lean mass of male mice at 22 weeks, measured by EchoMRI (n = 8/group). (H) gWAT weight in male mice at 24 weeks (n = 8/group). (I) Adipocyte diameter in gWAT in male mice (n = 8/group). (J) Representative F4/80 staining in male gWAT (10x, scale bars = 150□μm) (L) CLS quantified by F4/80 staining in male gWAT (n = 8/group; 3 fields per section). Data are represented as mean ± SEM. \**p* < 0.05, \*\**p* < 0.01, \*\*\**p* < 0.001, ns, not significant, by two-way ANOVA with Tukey’s multiple comparisons test (A-D, F-J, and L).

Consistent with these findings, HFD-fed female *Brca1*^+/−^ mice had significantly higher gonadal white adipose tissue (gWAT) weight (**Figure 2C**; WT: 2.09 ± 0.29 vs. *Brca1*^+/−^ : 2.80 ± 0.13, *p = 0*.*02*), while differences in inguinal WAT (iWAT) were not significantly different (**Figure S2C**). Histological analysis demonstrated larger adipocytes (**Figures 2D**; WT: 106.13μm ± 5.10 vs. *Brca1*^+/−^ : 120.95μm ± 3.43, *p = 0*.*02*) and increased crown-like structure (CLS) (**Figures 2E-2F**; WT: 2.07 ± 1.04 vs. *Brca1*^+/−^ : 6.41 ± 0.78, *p < 0*.*01*) in gWAT of female *Brca1*^+/−^ mice compared to WT.

In contrast, HFD-fed male *Brca1*^+/−^ mice had significantly lower fat mass than WT controls (**Figures 2G**; WT: 21.07g ± 0.57 vs. *Brca1*^+/−^: 18.24g ± 0.61, *p < 0*.*01*), with no difference in lean mass (**Figures 2H**). No differences in fat or lean mass were observed at earlier time points (10-12 weeks) (**Figures S2D and S2E**). gWAT weight, adipocyte diameter, or CLS were similar between genotypes (**Figures 2I-2L**), while iWAT weight was significantly lower in HFD-fed *Brca1*^+/−^ males (**Figure S2F**; WT: 3.08g ± 0.08 vs. *Brca1*^+/−^: 2.75g ± 0.10, *p = 0*.*02)*.

Adiposity can result from reduced energy expenditure, increased energy intake or both^14,15^. Since food intake was similar between genotypes in both sexes, we assessed energy expenditure using indirect calorimetry in female (10-11 weeks on diet) and male mice (12-13 weeks on diet). HFD-fed *Brca1*^+/−^ female mice exhibited significantly higher resting energy (REE) and total energy expenditure (TEE) in both light and dark cycles (**Figures S3A-S3F; (REE, light)** WT: 4.32 ± 0.15 kcal/day vs. *Brca1*^+/−^ : 4.76 kcal/day ± 0.11, *p = 0*.*03*, **(TEE, light)** WT: 5.06 kcal/day ± 0.15 vs. *Brca1*^+/−^ : 5.51kcal/day ± 0.11, *p = 0*.*02*),, (***REE, dark***) WT: 4.77 kcal/day ± 0.17 vs. *Brca1*^+/−^ : 5.25 kcal/day ± 0.12, *p = 0*.*03*), (**TEE, dark**) WT: 6.16 kcal/day ± 0.18 vs. *Brca1*^+/−^ : 6.66 kcal/day ± 0.11, *p = 0*.*02*) Covariate analysis was performed and differences in energy expenditure were no longer significant following correction for lean mass, while differences were not affected by correction for fat mass (**Table S1**). Male mice showed no genotype-dependent differences in energy expenditure (**Figures S3G-S3L**). Skeletal muscle weights measured at 22 weeks on diet showed no significant difference in either sex (**Figures S4A-S4D**). Moreover, no difference was obtained in muscle fiber composition in tibialis anterior (**Figures S4E-S4H**).

### 3. *Brca1* heterozygosity promotes high-fat diet-induced hepatic steatosis in both female and male mice

We next investigated whether *Brca1* heterozygosity affects hepatic lipid accumulation (steatosis) on a HFD. After 22 weeks of HFD feeding, female *Brca1*^+/−^ mice exhibited markedly increased hepatic steatosis compared with WT controls, as shown by H&E staining (**Figure 3A**). Histological scoring showed that steatosis and ballooning scores were higher in the female *Brca1*^+/−^ mice (**Figures 3B and 3C**), while lobular inflammation was similar between the groups (**Figure 3D**). Increased lipid accumulation was further confirmed by BODIPY staining (**Figure 3E**), with a significantly greater lipid droplet area per cell (**Figure 3F;** WT: 39.21μm^2^ ± 6.80 vs. *Brca1*^+/−^: 73.61μm^2^ ± 9.83, *p = 0*.*04*) and a trend toward larger droplet size (**Figure 3G**; WT: 10.01μm^2^ ± 1.91 vs. *Brca1*^+/−^: 17.27μm^2^ ± 2.59, *p = 0*.*07*).

**Figure 3.**
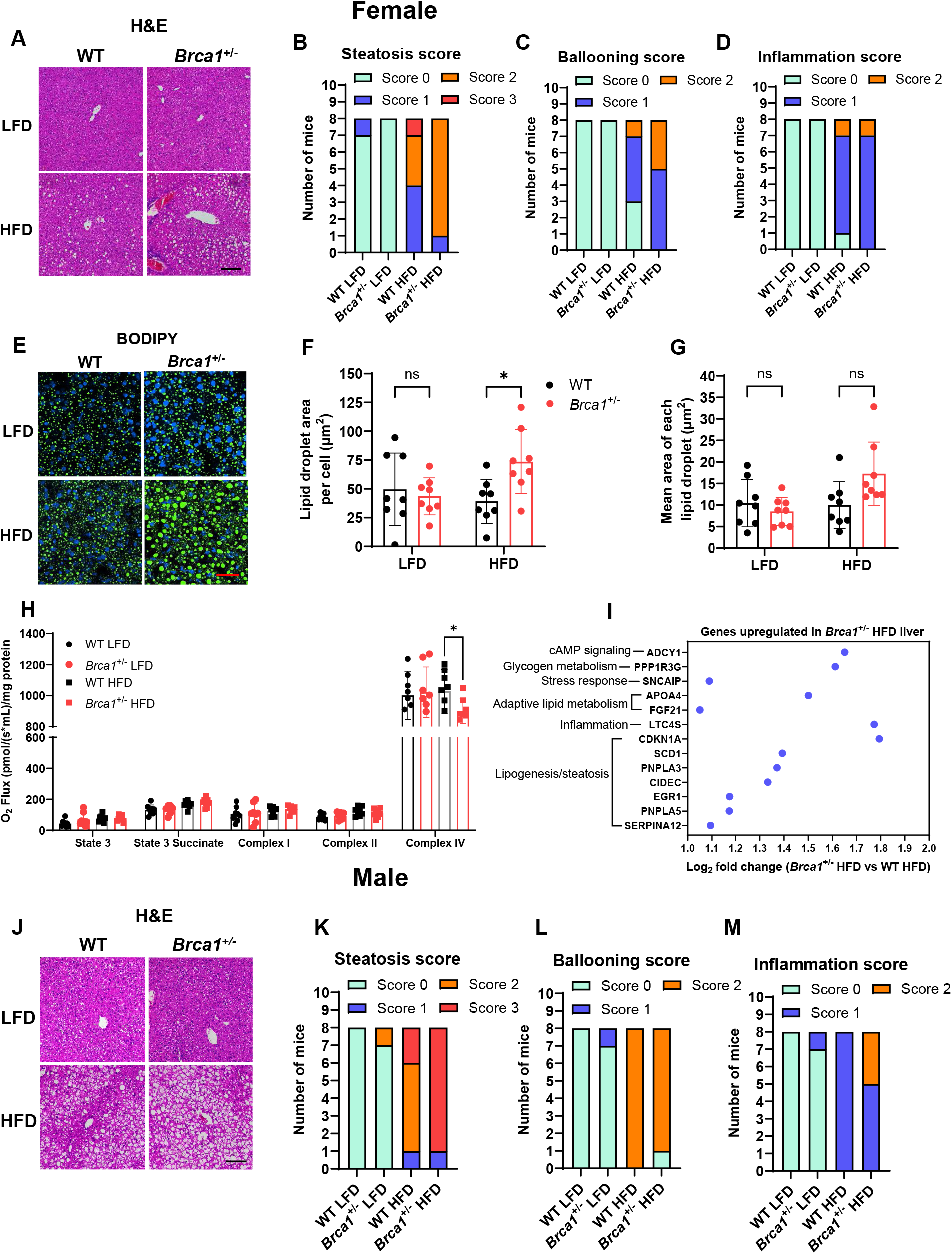
*Brca1* heterozygosity increases hepatic steatosis in both female and male mice with HFD. (A) Representative H&E-stained liver sections from WT and *Brca1*^+/−^ female mice fed LFD or HFD for 22 weeks (10x, scale bars = 150□μm) (B-D) Histology scoring of female liver sections for (B) steatosis, (C) hepatocyte ballooning, and (D) lobular inflammation (n = 8/group). (E) Representative immunofluorescence images of female liver sections stained with BODIPY (green) and Hoechst (nuclei, blue) (40x, scale bars = 40□μm, *n*□=□3 replicates). (F, G) Quantification of BODIPY staining in female livers showing (F) lipid droplet area per cell and (G) mean lipid droplet area (n = 8/group). (H) Mitochondrial respiration in isolated liver mitochondria of female mice: State 3, State 3 succinate, Complex I, Complex II, and Complex IV activities (n = 8/group). (I) Log_2_ fold change of selected upregulated genes in female Brca*1*^+/−^ HFD vs. WT HFD livers, grouped by functional annotation (n = 6/group). (J) Representative H&E-stained liver sections from male mice (10x, scale bars = 150□μm) (K-M) Histology scoring of male liver sections for (K) steatosis, (L) hepatocyte ballooning, and (M) lobular inflammation (n = 8/group). Data are represented as mean ± SEM. \**p* < 0.05, \*\**p* < 0.01, \*\*\**p* < 0.001, ns, not significant, by two-way ANOVA with Tukey’s multiple comparisons test (F, G, M, and N).

To explore potential metabolic alterations, respiration was assessed using Oroboros O2k high-resolution respirometers in isolated hepatic mitochondria. Mitochondrial complex IV activity was significantly reduced in HFD-fed female *Brca1*^+/−^ mice (**Figure 3H**; WT: 1027.44 ± 49.82 vs. *Brca1*^+/−^: 898.85 ± 28.23, *p = 0*.*03*), whereas no differences were observed in State 3 respiration, succinate-driven respiration, or complex I and complex II activities. RNA sequencing revealed upregulation of genes involved in lipid storage and steatosis (*Cdkna1, Scd1, Pnpla3, Cidec, EGR1, Pnpla5, Serpina12*), glycogen metabolism (*Ppp1r3g*), cAMP signaling (*Adcy1*), stress/inflammation (*Sncaip, Ltc4s*), and adaptive metabolic responses (*Fgf21* and *Apoa4)* (**Figure 3I**). HFD-fed *Brca1*^+/−^ male mice similarly showed increased hepatic steatosis compared with WT males (**Figure 3J and 3K**), with ballooning scores unchanged but greater lobular inflammation (**Figure 3L and 3M**).

### 4. Tirzepatide reduces body weight gain, improves glucose tolerance, and reduces hepatic steatosis in HFD-fed *Brca1* heterozygous knockout female mice

Given the heightened metabolic dysfunction in HFD-fed female *Brca1*^+/−^ mice, we tested whether tirzepatide, a dual GLP-1/GIP receptor agonist, could improve these phenotypes. Female *Brca1*^+/−^ mice received subcutaneous injection of tirzepatide or vehicle three times per week for 10 weeks (**Figure 4A**). Tirzepatide significantly reduced body weight compared with vehicle-treated HFD-fed *Brca1*^+/−^ mice (**Figure 4B;** HFD+VC: 37.28g ± 1.43 vs. HFD+TZP: 27.57g ± 1.13, *p* < *0*.*01*), accompanied by decreased food intake (**Figure 4C**). EchoMRI revealed that weight loss was primarily due to reduced fat mass as assessed at 10 weeks (**Figure 4D;** HFD+VC: 16.85g ± 1.37 vs. HFD+TZP: 8.16g ± 0.94, *p* < *0*.*01*), with only modest changes in lean mass (**Figure 4E;** HFD+VC: 19.82g ± 0.1934 vs. HFD+TZP: 18.53g ± 0.31, *p* < *0*.*01*).

**Figure 4.**
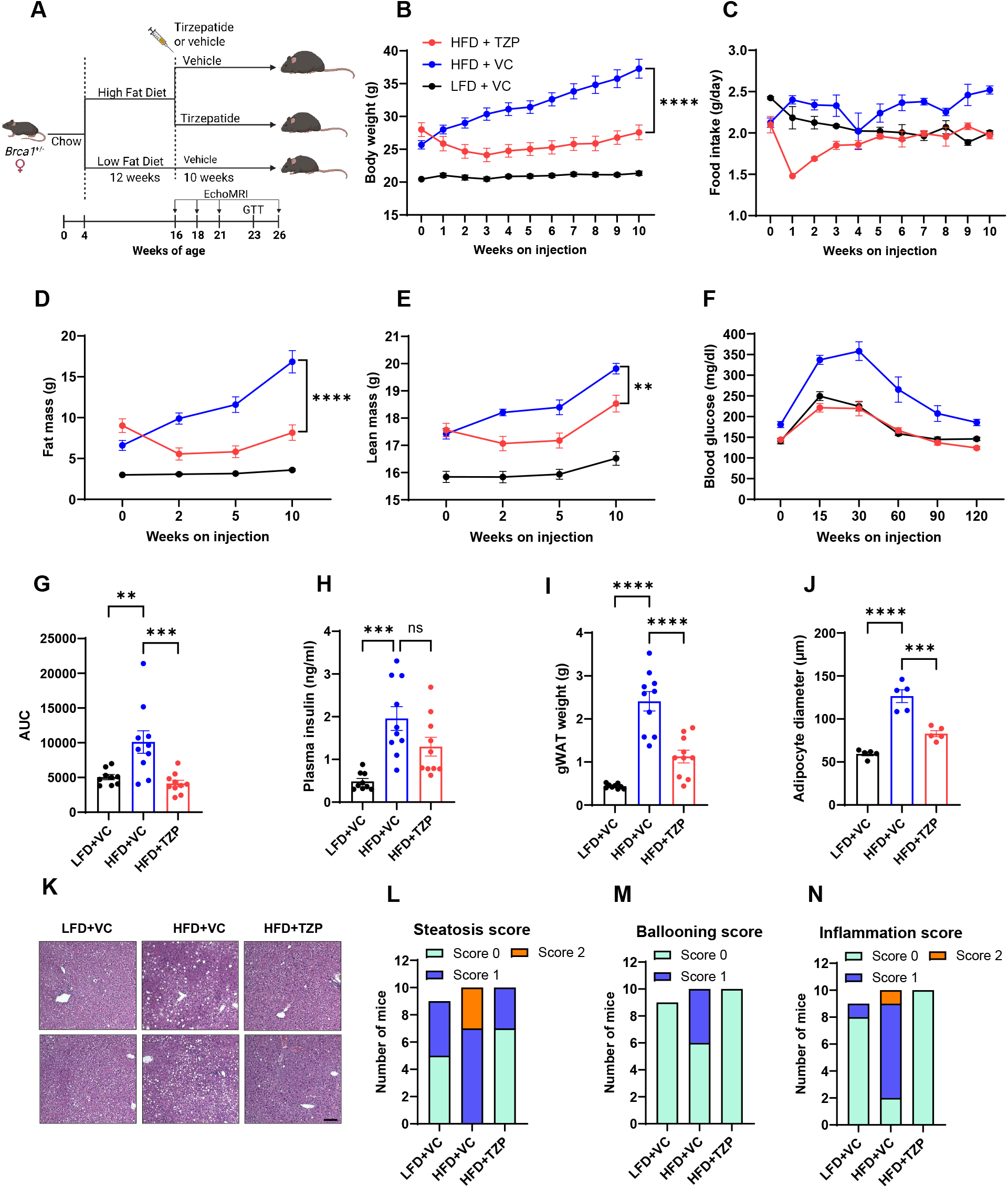
Tirzepatide improves adiposity, hepatic steatosis, and glucose tolerance in HFD-fed female *Brca1*^+/−^ mice. (A) Experimental design. (B) Body weight of *Brca1*^+/−^ female mice fed LFD or HFD and treated with either tirzepatide (TZP) or vehicle (VC) (n = 9-10/group). (C) Food intake measured over the treatment period (n = 10/per group). (D, E) (D) Fat mass and (E) lean mass progression assessed by EchoMRI. (F) Glucose tolerance test (GTT) and (G) area under the curve (AUC) after 7 weeks of treatment (n = 9-10/group). (H) Fasting plasma insulin levels after 10 weeks of treatment (2-hour fast, n = 9-10/group). (I) Gonadal white adipose tissue (gWAT) weight after 10 weeks of treatment (n = 9-10/group). (J) Adipocyte diameter in gWAT quantified from HE stained slides (n = 5/group). (K) Representative H&E-stained liver sections after 10 weeks of treatment (10x, scale bars = 100□μm). (L-N) Histological scoring for (L) steatosis, (M) hepatocyte ballooning, and (N) lobular inflammation (n = 9-10/group). Data are represented as mean ± SEM. \**p* < 0.05, \*\**p* < 0.01, \*\*\**p* < 0.001, ns, not significant, by one-way ANOVA with Tukey’s multiple comparisons test (B-J).

Glucose tolerance improved significantly in tirzepatide-treated mice (**Figures 4F and 4G**; AUC HFD+VC: 10107 ± 1616 vs. HFD+TZP: 4139 ± 455.10373.8, *p* < *0*.*01*). Fasting plasma insulin levels tended to be lower in tirzepatide-treated mice (**Figure 4H**; HFD+VC: 1.96 ± 0.28ng/ml vs. HFD+TZP: 1.30ng/ml ± 0.22, *p = 0*.*09*). Adipose tissue was also significantly affected in tirzepatide-treated mice, as evidenced by decreased gWAT weight (**Figures 4I**; HFD+VC: 2.41g ± 0.23 vs. HFD+TZP: 1.13g ± 0.15, *p* < *0*.*01*) and smaller adipocytes (**Figure 4J**; HFD+VC: 126.50μm ± 7.40 vs. HFD+TZP: 82.92μm ± 3.50, *p* < *0*.*01*). Tirzepatide also ameliorated hepatic steatosis, reducing steatosis score, hepatocyte ballooning, and lobular inflammation compared with vehicle-treated mice (**Figures 4K-4N**).

## DISCUSSION

In this study, we demonstrate that heterozygous loss of *Brca1* alters metabolic adaptation to HFD feeding in a sex-dependent manner. Compared with WT female mice, *Brca1*^+/−^ female mice developed exacerbated HFD-induced obesity, glucose intolerance, adipose tissue inflammation, and severe hepatic steatosis. This is a profound finding given that female mice are typically protected from HFD-induced steatosis. In contrast, compared with WT male mice, *Brca1*^+/−^ male mice were relatively protected from systemic metabolic dysfunction, despite developing hepatic steatosis. These findings expand the functional scope of Brca1 beyond genome maintenance and identify a previously unrecognized role ofBrca1 in hepatic lipid homeostasis.

The observation that hepatic steatosis occurs in both sexes suggests a liver-intrinsic vulnerability associated with *Brca1* heterozygosity. Mitochondrial oxidative metabolism plays a central role in maintaining lipid homeostasis by enabling efficient fatty acid oxidation and preventing excessive lipid storage^16,17^. Accordingly, a reduction in mitochondrial respiratory capacity may therefore limit the liver’s ability to appropriately process increased lipid influx during high-fat feeding, predisposing hepatocytes to triglyceride accumulation. In parallel, the observed transcriptional profile suggests a shift toward metabolic programs that favor nutrient storage and adaptive signaling rather than oxidative lipid disposal. Such reprogramming may initially serve as a compensatory response to nutrient excess, aimed at buffering metabolic stress; however, in the context of impaired mitochondrial function, these adaptations may ultimately exacerbate lipid accumulation and contribute to steatosis.

Sexual dimorphism in systemic metabolic outcomes was a prominent feature of *Brca1* heterozygosity under HFD feeding. Our findings reveal clear sex-dependent effects of *Brca1* heterozygosity on metabolic adaptation, with females exhibiting heightened systemic metabolic vulnerability compared with WT controls, while males were relatively protected from overt systemic dysfunction. Consistent with these observations, Hembruff *et al*. reported sex-specific differences in body composition, energy expenditure, and glucose handling in humanized *BRCA1* knock-in mice, although these phenotypes were assessed at 6 months and 1 year of age without dietary intervention^18^. Despite differences in genetic models, our study extends this work by demonstrating that *Brca1* plays an important role in regulating metabolic health, in a diet-dependent and sex-dependent manner. Given that female mice are typically resistant to HFD-induced hepatic steatosis, the loss of this protection in *Brca1*^+/−^ females underscores the importance of Brca1 as a biological variable in metabolic regulation in the liver. The worsening of hepatic steatosis in *Brca1*^+/−^ males, without significant systemic metabolic alterations, highlight sex-independent effects of Brca1 intrinsic to liver metabolism.

*Brca1* heterozygosity also induced HFD-induced increases in lean mass in females, an effect not observed in male mice. Given that lean mass is a major determinant of resting energy expenditure^19,20^, this increase likely contributes to the elevated resting energy expenditure observed in HFD-fed female *Brca1*^+/−^ mice. Consistent with this interpretation, the genotype-dependent difference in resting energy expenditure was no longer significant after adjustment for lean mass. Despite the increase in lean mass, direct assessment of skeletal muscle mass, including gastrocnemius and tibialis anterior, revealed no significant differences between genotypes, nor were there differences observed in tibialis anterior muscle fiber composition. These findings suggest that other lean tissue compartments may contribute to the elevated lean mass and energy expenditure in female *Brca1*^+/−^ mice. In contrast to our findings, Tarpey *et al*. reported that skeletal muscle-specific deletion of *Brca1* reduced the proportion of type IIb fibers in the tibialis anterior^21^. This difference likely reflects the use of a systemic *Brca1* heterozygous model in our study, rather than complete muscle-specific deletion in the knockout model.

Finally, the efficacy of tirzepatide in reversing metabolic dysfunction in *Brca1*^+/−^ females demonstrates that *Brca1*-associated metabolic defects are amenable to pharmacological intervention. These findings suggest that incretin-based therapies may mitigate the risk of metabolic disease in *BRCA1* mutation carriers exposed to obesogenic environments.

In summary, *Brca1* heterozygosity predisposes mice to diet-induced metabolic dysfunction in a sex-dependent manner, with females exhibiting heightened systemic metabolic vulnerability relative to WT controls, while both sexes develop hepatic steatosis. This study highlights a clinically relevant metabolic liability that may be underappreciated, particularly given the limited epidemiological data on metabolic outcomes in male *BRCA1* mutation carriers. It also demonstrates the potential of incretin-based therapies, to mitigate metabolic risk in *BRCA1* mutation carriers. Given our prior studies elucidating the role of obesity and poor metabolic health in driving cancer risk in this population^10^, these findings would support further evaluation and monitoring of liver health in carriers.

### Limitations of the study

This study used male and female *Brca1* heterozygous knockout mice to assess the impact of a high-fat diet on metabolic health. Although designed to mimic one of the most prevalent pathogenic germline *BRCA1* heterozygous mutations, validation in human cohorts will be necessary to determine the relevance of *BRCA1* heterozygosity to hepatic steatosis and metabolic dysfunction.

## RESOURCE AVAILABILITY

### Lead contact

Inquiries regarding resources, reagents, and further information should be directed to, and will be fulfilled by, the lead contact, Dr. Kristy A. Brown.

### Material availability

This study did not generate new unique reagents. Reagents are available upon request from the lead contact.

## Data availability

- Raw data underlying all graphs can be found in Data S1.
- Any additional information is available from the lead contact upon request.

## ACKNOWLEDGMENTS

We thank Katherine McCartney for mouse husbandry and Dr. Espen Spangenburg (East Carolina University) for skeletal muscle fiber analysis methodology. Schematics were created using BioRender. We thank all the members of the Brown laboratories for helpful discussion. This work was supported by funding from the University of Kansas Cancer Center (NIH P30CA168524), Kansas Center for Metabolism & Obesity Research Center of Biomedical Research Excellence NIH P20GM144269 and 3P20GM144269-03S1, NIH R01CA215797, V Foundation for Cancer Research (T2025-007) to K.A.B. and the University of Kansas Biomedical Research Training Program Fellowship (NIH P20GM103418) to S.P. Histological work and use of the Nikon CSU-W1 SoRa Spinning-Disk Super-Resolution Confocal microscope at the University of Kansas Medical Center Integrated Imaging Core were supported by NIH U54 HD 090216.

## AUTHOR CONTRIBUTIONS

K.A.B., J.P.T., and S.P conceived and designed the study. S.P., L.Q., C.K., P.B, X.L, and C.S.M. performed the experiments and analyzed the data. P.D. analyzed the sectioned slides and interpreted the histological data. O.P. and H.M.T. performed GSIS experiments and analyzed the data. D.C.K. reviewed and validated the statistical analyses. K.A.B. and S.P. wrote the manuscript.

## DECLARATION OF INTERESTS

The authors declare no competing interests.

## STAR METHODS

### KEY RESOURCES TABLE

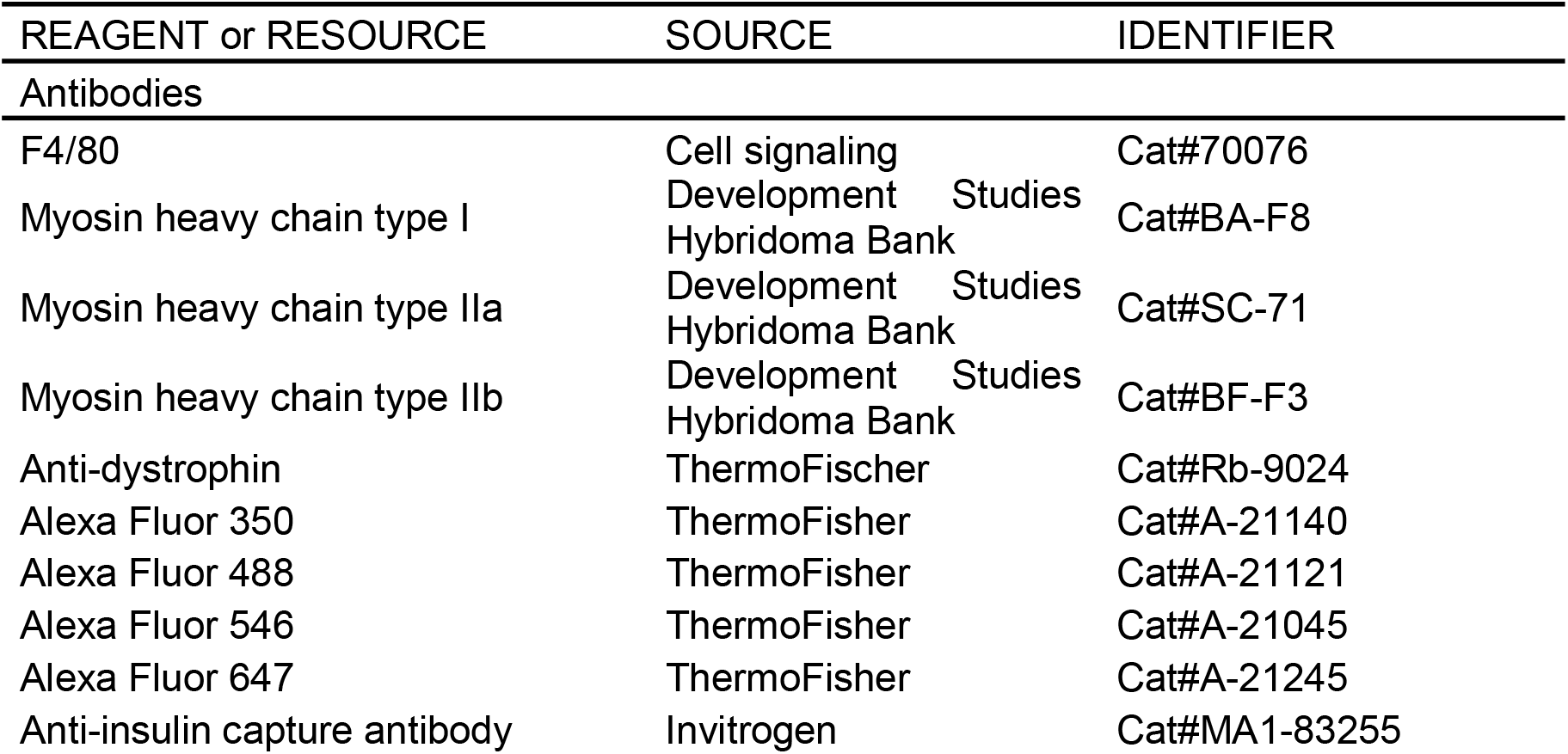
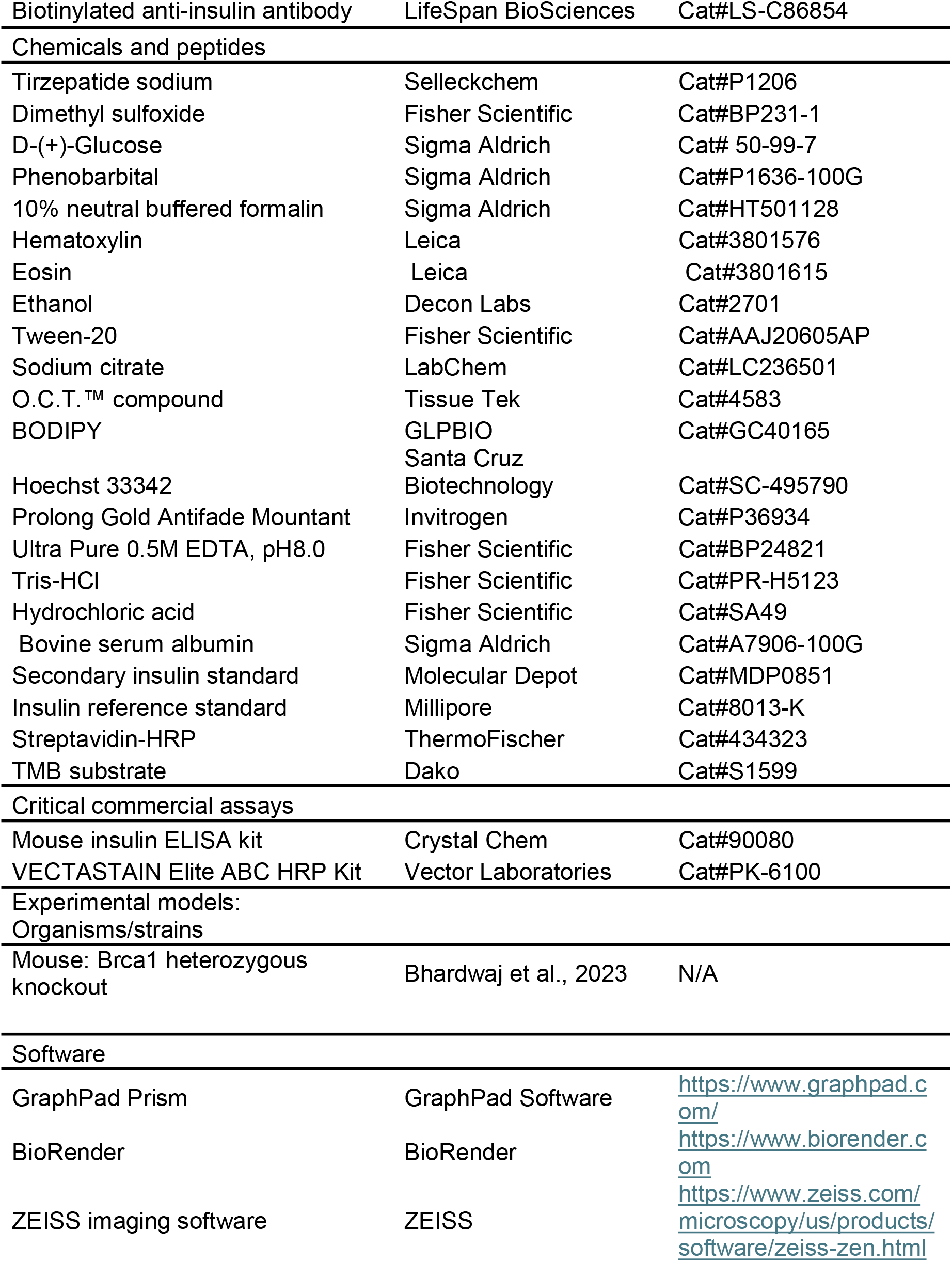
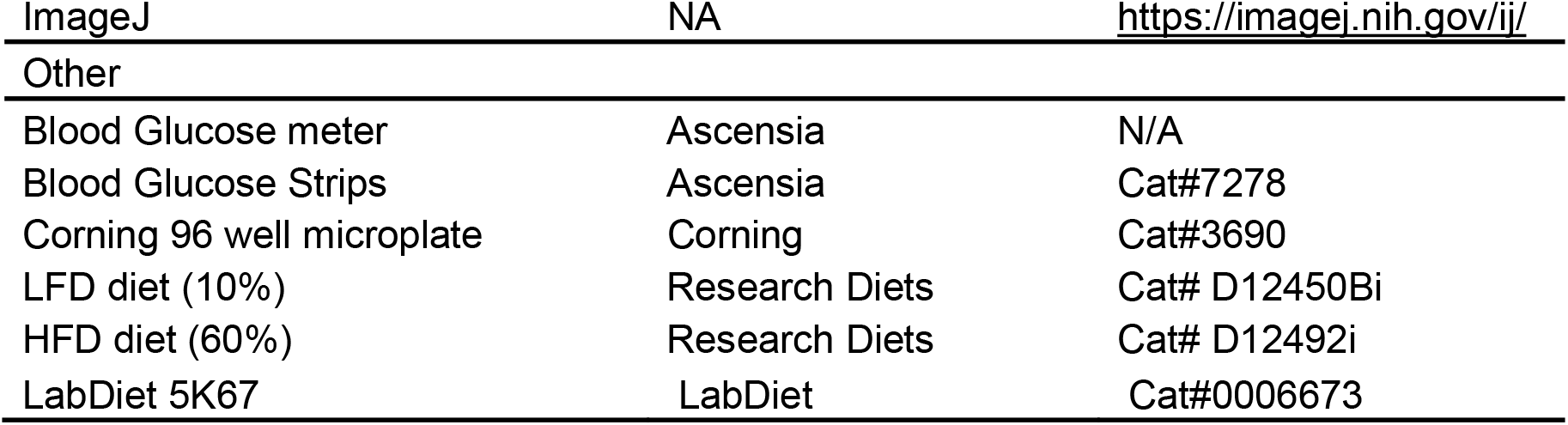

## EXPERIMENTAL MODEL

### Mouse models

The animal protocol was approved by the Institutional Animal Care and Use Committee at the University of Kansas Medical Center (animal protocol number: IPROTO2023-306). Mice were anesthetized with phenobarbital (200 mg/kg) prior to the terminal procedures following a 2-hour fast in the light cycle (8:00-10:00). The generation of *Brca1*^+/−^ mice has been previously described^10^. For the present study, male and female *Brca1*^+/−^ mice on a C57BL/6 background were bred together to generate *Brca1*^+/−^ and WT littermates. Genotyping was performed by Transnetyx Inc. (Cordova, TN, USA) using a real-time PCR-based TaqMan^®^ assay. Tail snips were collected and submitted according to the company’s standard protocol. Mice were housed on a 12-h light/dark cycle (lights on 6:00, off at 18:00) with ad libitum access to water and food (LabDiet 5K67). At 4 weeks of age, female *Brca1*^+/−^ and WT mice were fed either a 60 kcal% HFD (D12492i, Research Diets) or a 10 kcal% LFD (D12450Bi, Research Diets) ad libitum for 22 weeks until euthanasia, whereas male mice were fed the same diets for 24 weeks. Body weight and food intake were measured weekly. Energy intake was calculated by measuring # of g diet consumed per mouse x 3.85kcal/g for LFD or 5.24kcal/g for HFD. In a separate cohort, female *Brca1*^+/−^ mice were fed either a HFD or LFD for 12 weeks beginning at 4 weeks of age. Mice were euthanized for tissue collection at the end of the dietary intervention.

## METHOD DETAILS

### Tirzepatide injection

At 4 weeks of age, female *Brca1*^+/−^ mice were randomized to receive either HFD (n=20) or LFD (n=10). After 12 weeks on the diet, HFD-fed mice were further randomized to receive subcutaneous injection of either tirzepatide (TZP, 10 nmol/kg; n=10) or vehicle (5% DMSO, 95% 40 mM Tris HCl, pH 8.0, n=10). Control mice continued on the LFD and received the vehicle injection. Subcutaneous injections were performed three times a week (Monday, Wednesday, and Friday) and continued for 10 weeks. Mice were euthanized for tissue collection at the end of the injection period.

### Body composition analysis

Body composition was determined longitudinally utilizing magnetic resonance via the EchoMRI-900 analyzer (EchoMRI, Houston, TX, USA) at predetermined intervals. The difference between total body mass and fat mass was used to calculate fat free mass in all animals.

### Indirect calorimetry

Indirect Calorimetry analysis was performed at the University of Kansas Medical Center Metabolic Obesity Phenotyping Facility using a Promethion continuous indirect calorimetry system (Sable Systems International, Las Vegas, NV) as previously described^15^. Since indirect calorimetry requires mice to be singly housed, all mice (both male and female) were individually housed one week prior to the experiment. Macros available from the manufacturer were used to calculate total energy expenditure (TEE) based on the Weir equations and respiratory quotient (RQ) by VCO_2_/VO_2_. To calculate resting energy expenditure (REE), the Promethion designed macro pulled out the lowest 30 minutes period of EE (light cycle) which was then extrapolated to a 24-hour period. The difference between TEE and REE was used to determine non-resting energy expenditure (NREE). Mice were given 2 days to acclimatize to new cages before data was recorded. At the end of the indirect calorimetry, female mice were regrouped as previously housed, while male mice remained singly housed until euthanasia.

### Glucose tolerance test

For glucose tolerance tests, mice were fasted for 6□hours and intraperitoneally (i.p.) injected with glucose (1□g/kg body weight). Blood was taken by tail vein puncture, and glucose levels were measured using glucometer (Ascensia) at indicated time points.

### Insulin ELISA from murine plasma

Blood was collected by terminal cardiac puncture using a syringe rinsed with EDTA (0.5 M), transferred to Eppendorf tubes, and kept on ice. Mice were fasted for 2 h before blood collection. Samples were centrifuged at 1,000 x g for 15 minutes at 4°C to separate plasma. Plasma samples were stored at −80 °C until use, with freeze-thaws kept to a minimum. Plasma insulin concentrations were measured using a mouse insulin ELISA kit (Crystal Chem, cat. #90080) according to the manufacturer’s instructions. Samples and standards were assayed in duplicate.

### Glucose-stimulated insulin secretion (GSIS)

GSIS was performed as previously described^22^. In short, islets were prepared from pancreata by CIzyme RI digestion (VitaCyte, #005-1030), handpicked, and then incubated in a humidified atmosphere in RPMI 1640 culture medium (Gibco, 21875091) supplemented with 10% (vol/vol) FBS, 100 U/ml penicillin, 100 U/ml streptomycin, 20 mM HEPES, 2mM L-glutamine, 50 μm 2-Mercaptoethanol, and 0.5% BSA. Freshly isolated islets were incubated overnight before the experiment. To determine insulin secretion, on the day of the experiment, 10 size-matched islets were picked for each group and pre-incubated for 2 h at 37□°C in Krebs-Henseleit Ringer Bicarbonate Buffer (KRB buffer) composed of 118.41□mM NaCl, 4.69□mM KCl, 1.18□mM MgSO_4_·7H_2_O, 1.18 mM KH_2_PO_4_, 25 mM NaHCO_3_, 5□mM HEPES, 2.52□mM CaCl_2_.2H_2_O, 0.5% bovine serum albumin, and supplemented with 1.7□mM glucose. Supernatant was aspirated without disturbing the islets and 200 μl of fresh KRB supplemented with 1.7 mM glucose was added at 37°C for 1 h. Supernatant was collected and 200 μl of fresh KRB supplemented with 16.7 mM glucose was added at 37°C for 1 h.

Insulin concentrations in the supernatants were measured using a sandwich ELISA. Plates were coated overnight at room temperature with anti-insulin capture antibody (Invitrogen, #MA1-83255) and blocked with 5% BSA for 1 hour at room temperature. Sample supernatants and a secondary insulin standard (Molecular Depot, #MDP0851), calibrated against the reference standard (Millipore, #8013-K), were incubated overnight at 4°C. Bound insulin was detected using a biotinylated anti-insulin antibody (LifeSpan BioSciences, #LS-C86854) for 2 hours at room temperature, followed by streptavidin-HRP (ThermoFischer, #434323) for 30 minutes. Signal was developed using TMB substrate (Dako, #S1599), and the reaction was stopped with 0.18 M H_2_SO_4_. Absorbance was measured at 450 nm using a Synergy H1 microplate reader (BioTek).

### Histology and immunohistochemistry staining

Mouse tissues (liver, gWAT, and iWAT) were fixed in 10% neutral buffered formalin (Sigma Aldrich, HT501128) at 4°C for 24 h. Then, tissues were rinsed in PBS and stored in 70% ethanol until use. Fixed tissues were embedded in paraffin, sectioned 5 μm thick, and stained with Hematoxylin and Eosin solution (H&E). H&E-stained slides were pictured using a Revolve microscope. For immunohistochemistry, formalin-fixed and paraffin-embedded adipose tissue sections were deparaffinized and rehydrated prior to antigen unmasking by boiling in 10 mM sodium citrate, pH 6.0, with 0.05% Tween-20 for 30 minutes. Sections were blocked in normal serum and incubated with F4/80 antibody (Cell signaling #70076) overnight. Endogenous HRP activity was quenched with incubation with 3% hydrogen peroxide for 10 minutes. Secondary antibody staining was performed using the Vectastain ABC kit (Vector Laboratories) and detected with 3,3′-diaminobenzidine (DAB). Sections were counterstained with hematoxylin prior to dehydration and coverslip placement.

The liver histology was evaluated by an expert liver pathologist who was blinded to the different treatments. The amount of steatosis (percentage of hepatocytes containing fat droplets) was scored as 0 (<5%), 1 (5–33%), 2 (34–66%), and 3 (>66%). Hepatocyte ballooning was classified as 0 (none), 1 (few), or 2 (many cells/prominent ballooning). Foci of lobular inflammation were scored as 0 (no foci), 1 (1–2 foci per 20× field), 2 (2–4 foci per 20× field), and 3 (>4 foci per 20× field). Images were acquired using a Leica DME microscope and Leica ICC50HD camera (Leica, Wetzlar, Germany) and analyzed using Leica LAS EZ software.

H&E-stained slides of gWAT and iWAT were imaged using a Revolve microscope at 10x magnification. Adipocyte diameters were measured using ZEISS imaging software. For each image, a straight line was drawn across the field, and the diameters of fifteen adipocytes intersected by the line were recorded. Measurements were performed in two fields of view per sample.

### BODIPY staining liver

Liver samples were collected, placed in cryomolds, covered with O.C.T.™ compound (Tissue Tek, #4583) and frozen on dry ice-chilled isopentane. Frozen tissue sections were cut 8 μM thick and mounted on slides. For BODIPY staining, frozen tissue sections were incubated with 25□μM BODIPY Reagent (GLPBIO, GC40165, Montclair, CA) for 30□minutes at 37□°C in the dark. Then the sections were rinsed with PBS for 5 minutes and fixed at room temperature with 10% neutral-buffered formalin for 20□minutes. After washing with PBS for 5 minutes, the slides were counterstained with Hoechst 33342 nuclear stain (Santa Cruz Biotechnology, #SC-495790) for 5 minutes at 1:1000 dilution. The slides were washed with PBS for 5 minutes and mounted with Prolong Gold Antifade Mountant (Invitrogen, #P36934). Lipid droplets were imaged by using the Nikon CSU-W1 Spinning-disk confocal 60x objective and an intermediate 4x magnification changer with SoRa module. BODIPY fluorescence intensities were quantified using ImageJ.

### Liver Mitochondrial Isolation and Respiration

Liver mitochondrial were isolated as previously described^23-25^. Briefly, livers were rapidly excised and immediately submerged in 8 mL of ice-cold mitochondrial isolation buffer (220 mM mannitol, 70 mM sucrose, 10 mM Tris, 1 mM EDTA, pH 7.4). Tissue was homogenized on ice with a Teflon pestle. Homogenates were centrifuged at 1,500 × g for 10 minutes at 4 °C, strained through gauze, and the resulting supernatant was centrifuged at 8,000 × g for 10 minutes at 4 °C. The supernatant was discarded, and the mitochondrial pellet was resuspended in 6 mL of isolation buffer using 3–4 passes of glass-on-glass homogenization, followed by centrifugation at 6,000 × g for 10 minutes at 4 °C. This washing step was repeated using 4 mL of isolation buffer supplemented with 0.1% fatty acid–free BSA and lower centrifugation speed (4 °C, 10 minutes, 6000*g*). The final mitochondrial pellet was resuspended in 300 to 500 µL modified MiR05 mitochondrial respiration buffer (0.5 mM EGTA, 3 mM MgCl_2_, 60 mM KMES, 20 mM glucose, 10 mM KH_2_PO_4_, 20 mM HEPES, 110 mM sucrose, 0.1% BSA, pH 7.1) and used in mitochondrial respiration studies.

Mitochondrial oxygen consumption (JO_2_; pmol·s^−1^·mL^−1^) was measured using an Oroboros O2k high-resolution respirometer (Oroboros Instruments, Innsbruck, Austria) as previously described^23-25^. Following air calibration, isolated mitochondria were added to chambers containing 2 mL of modified MiR05 respiration buffer maintained at 37 °C and continuously stirred at 750 rpm. Respiration buffer was supplemented with 2 mM malate, 63.5 μM free CoA, and 2.5 mM L-carnitine. Glutamate-supported respiration was initiated with 2 mM glutamate, followed sequentially by the addition of 2.5 mM ADP (State 3), 10 mM succinate (state 3S), 2 μM rotenone (complex I inhibitor), 5 mM malonate (complex II inhibitor), and 2 mM ascorbate + 0.5 mM TMPD (complex IV substrate). Complex I activity was calculated as the difference between State 3S respiration and respiration following rotenone addition, whereas complex II activity was calculated as the difference between respiration following rotenone and malonate addition. Data were acquired and analyzed with DatLab software (version 7, Oroboros Instruments) and normalized to total mitochondrial protein content in each chamber.

### Muscle fiber type

Muscle fiber type was measured in the tibialis anterior (TA) muscle as a result of mixed muscle fiber type, as described previously^26^. Images of muscle cross-sections were taken by a blinded investigator from six different locations across the belly of the muscle section and were probed with primary antibodies against myosin heavy chain type I (#BA-F8), IIa (#SC-71) and IIb (#BF-F3) and anti-dystrophin (#Rb-9024). Myosin heavy chain types I, IIa and IIb and dystrophin were stained with secondary fluorescence probes 350, 488, 546 and 647, respectively. Stained cross-sections were imaged using the Nikon CSU-W1 Spinning-disk confocal 60x objective and an intermediate 4x magnification changer with SoRa module. Fiber types were assessed using Image J (NIH, Bethesda, MD, USA), as described previously^26^. All images were coded and randomized to allow for blinded analyses.

### RNA sequencing and bioinformatic analysis

Total RNA was isolated from 20–50□mg of snap-frozen liver tissue using QIAshredder columns (Qiagen, #79656) and the Rneasy® Mini Kit (Qiagen, #74106) according to the manufacturer’s instruction. The integrity and quality of the extracted RNA were assessed using an Agilent TapeStation 5400 System (hagilent.com).

Library preparation was performed by Novogene (Sacremento, CA, USA) using a non-strand-specific mRNA library preparation protocol. Briefly, mRNA was purified from total RNA using poly-T oligo-attached magnetic beads and fragmented. First-strand cDNA was synthesized using random hexamer primers, followed by the second-strand cDNA synthesis. The resulting double-stranded cDNA underwent end repair, A-tailing, adapter ligation, size selection, PCR amplification, and purification. Library quality and concentration were assessed using Qubit fluorometer, real-time PCR, and an Agilent Bioanalyzer for fragment size distribution. After quality control, libraries were pooled based on effective concentration and the targeted data amount and subjected to 150-bp paired-end sequencing on the NovaSeq X Plus platform (Illumina).

RNA sequences were mapped to Mus musculus GRCh39/mm39 reference genome using HISAT2 (2.2.1). FeatureCounts (2.0.6) was used to count the read numbers mapped to each gene. Differential expression analysis was performed using the DESeq2 R package (1.42.0).

## QUANTIFICATION AND STATISTICAL ANALYSIS

Data are presented as the mean ± SEM as described in the figure legend. Statistical analyses were performed with GraphPad Prism 10 and SPSS (IBM SPSS Statistics).

One-way ANOVA was used for comparisons among three groups. Two-way ANOVA was used for group comparisons involving two or more factors, with repeated measures applied where appropriate. For energy expenditure analysis, analysis of covariance (ANCOVA) was performed in SPSS, with genotype as fixed effect, and body weight, fat mass, or lean mass included as covariates in separate models. Outliers were identified and removed using Grubb’s test. The value of *P*□<□0.05 was considered significant. Statistically significant differences are shown with asterisks (\**P*□<□0.05, \*\**P*□<□0.01, \*\*\**P*□<□0.001). The statistical details of experiments can be found in the figures and figure legends.

## Notes

### Competing Interest Statement

The authors have declared no competing interest.

