## Supplementary Figures for "*Brca1* heterozygosity leads to hepatic steatosis in male and female mice despite sexually dimorphic effects on systemic metabolism"

### Supplemental Figure 1

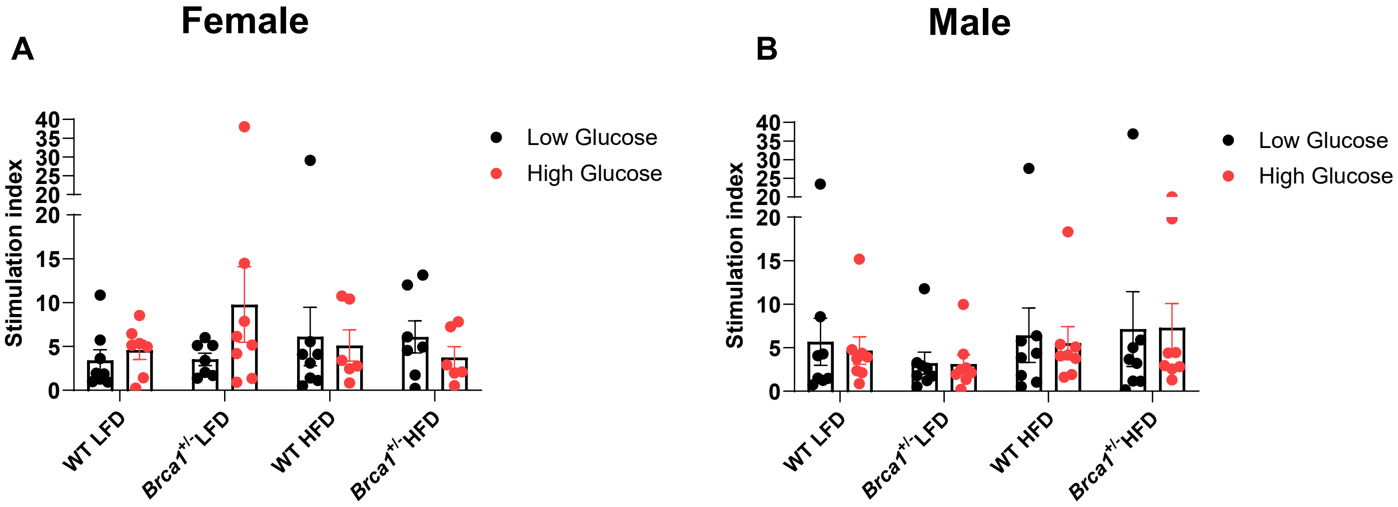


**Supplementary Figure 1: Glucose-stimulated insulin secretion (GSIS) in female and male mice.** GSIS was performed in islets of female (A) and male (B) mice after 22 weeks on diet (n = 8 per group) using low glucose (1.7 mM) or high glucose (16.7 mM). Data are represented as mean ± SEM. *$p < 0.05$ by two-way ANOVA with Tukey's multiple comparisons test.

### Supplemental Figure 2

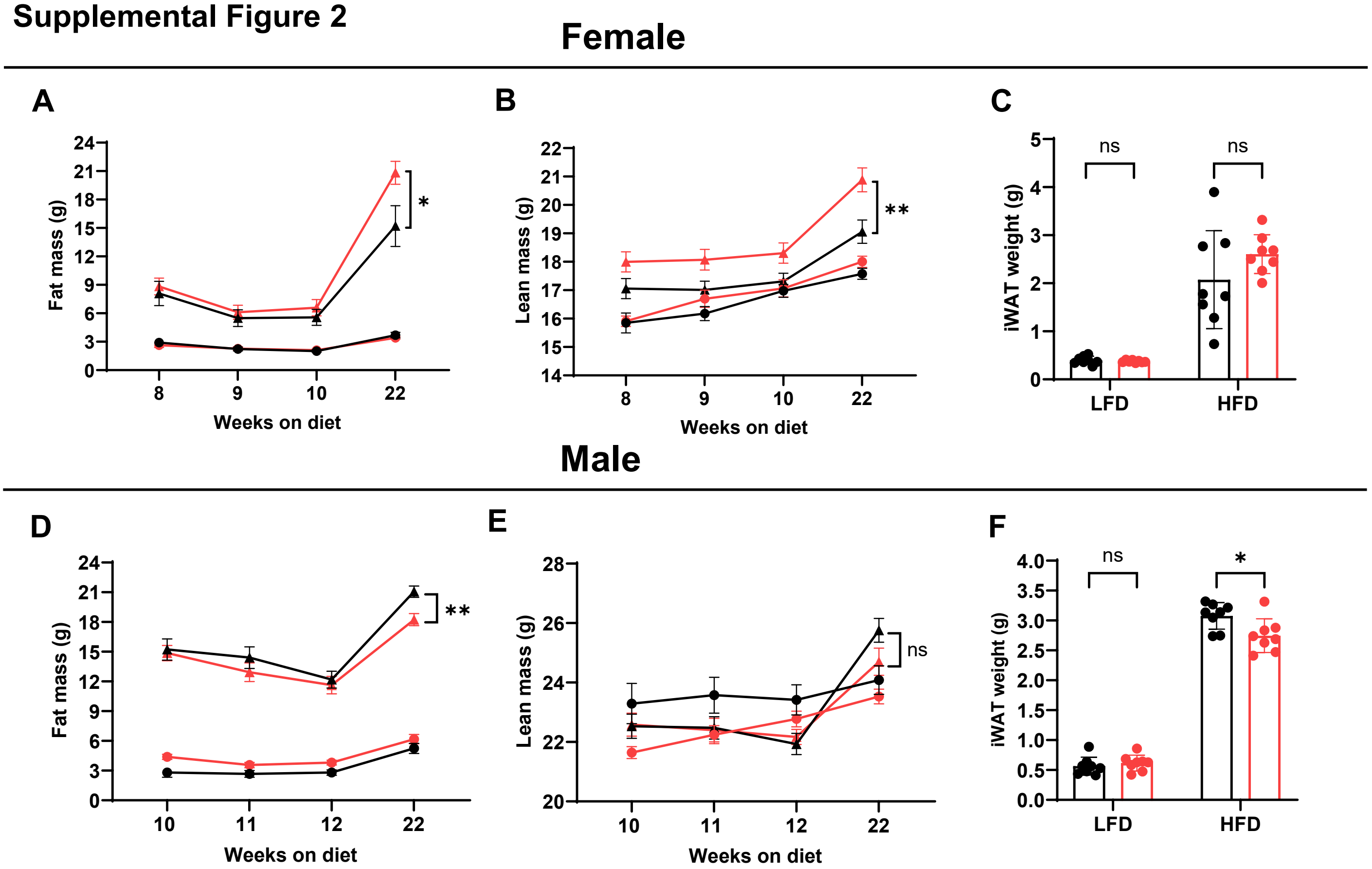


**Supplementary Figure 2: Fat mass, lean mass, and iWAT weight of female and male mice.** (A) Fat mass and (B) lean mass progression over the 22-week dietary period, assessed by EchoMRI. (C) Inguinal white adipose tissue (iWAT) weight of female mice after 22 weeks on diet (n = 8 per group). (D) Fat mass and (E) lean mass progression over the 22-week dietary period, assessed by EchoMRI. (F) iWAT weight of male mice after 24 weeks on diet (n = 8 per group). Data are represented as mean ± SEM. *$p < 0.05$, ns, not significant, by two-way ANOVA with Tukey's multiple comparisons test (A-F).

**Supplemental Figure 3**

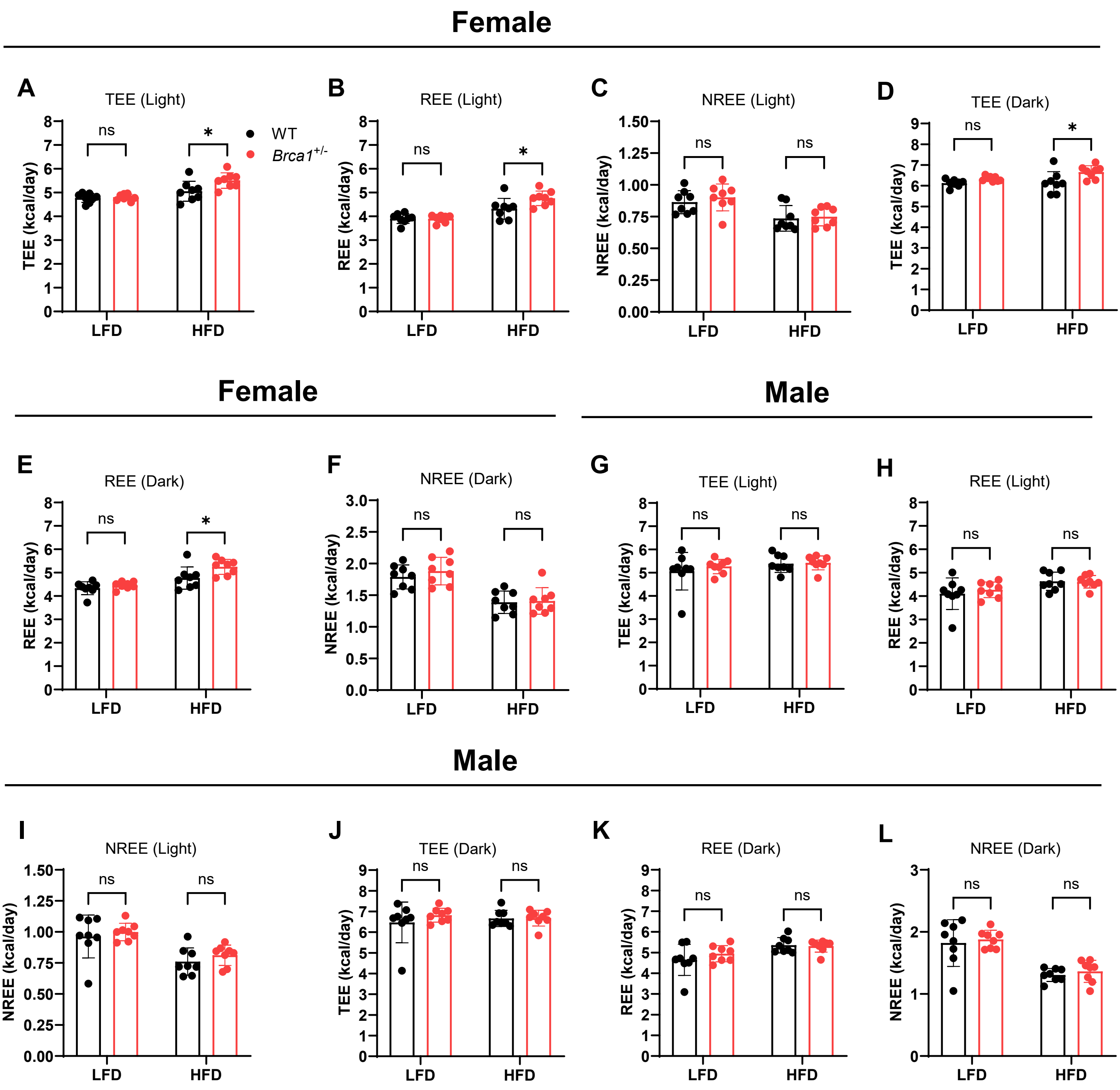


**Supplementary Figure 3: Female Brca1 heterozygous knockout mice have higher total (TEE) and resting energy expenditure (REE), but no difference in male mice.** Indirect calorimetry was performed when female mice were 10-11 weeks on diet. (A-C) (A) TEE, (B) REE, and (C) NREE during light cycle. (D-F) (D) TEE, (E) REE, and (F) NREE during dark cycle.(n = 8 per group). Indirect calorimetry was performed when male mice were 12-13 weeks on diet. (G-I) (G) TEE, (H) REE, and (I) NREE during light cycle. (J-L) (J) TEE, (K) REE, and (L) NREE during dark cycle.(n = 8 per group) ). Data are represented as mean ± SEM. **p* $< 0.05$, ns, not significant, by two-way ANOVA with Tukey's multiple comparisons test (A-P).

**Supplemental Figure 4**

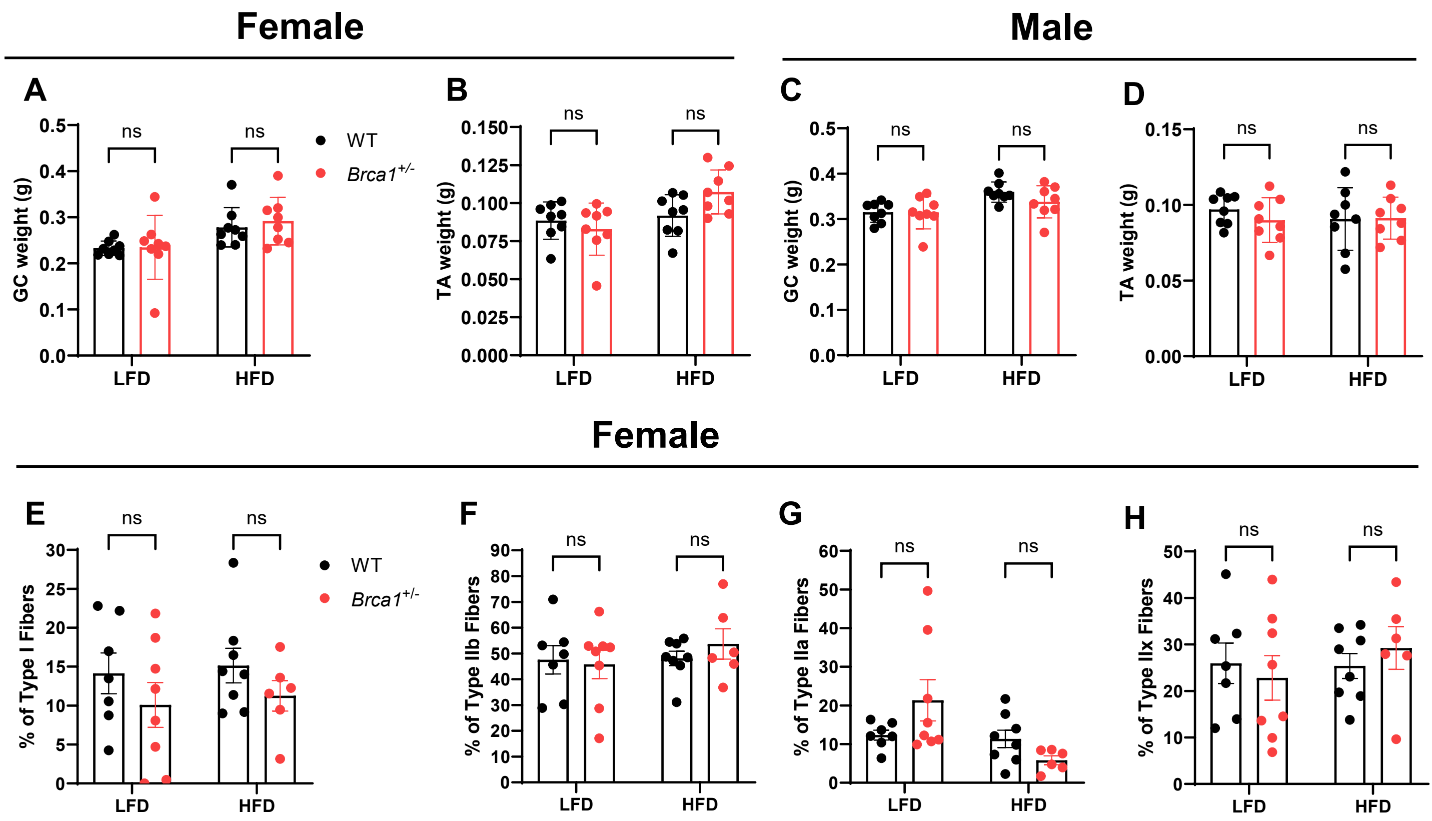


**Supplementary Figure 4: Gastrocnemius and tibialis anterior muscles weight in female and male mice and fiber type composition in female tibialis anterior muscle.** (A) Gastrocnemius (GC) and (B) tibialis anterior (TA) muscle wt of female and (C, D) male mice. (E-H) % type I, type IIb, type IIa, and type IIx fibers in TA muscle of female mice (n = 6-8 per group) ). Data are represented as mean ± SEM. $*p < 0.05$, ns, not significant, by two-way ANOVA with Tukey's multiple comparisons test (A-P).

**Supplemental Table 1**

**Table 1.** ANCOVA analysis between WT HFD vs *Brca1*$^{+/-}$ HFD (female)

| **Group** | **Parameter** | **Adjusted to body wt (*P* value)** | **Adjusted to fat mass (*P* value)** | **Adjusted to lean mass (*P* value)** |
|---|---|---|---|---|
| WT HFD vs *Brca1*$^{+/-}$ HFD | TEE (Light) | 0.076 | 0.016* | 0.394 |
| | TEE (Dark) | 0.115 | 0.040* | 0.40 |
| | REE (Light) | 0.148 | 0.032* | 0.563 |
| | REE (Dark) | 0.226 | 0.075 | 0.483 |
| | NREE (Light) | 1.0 | 1.0 | 1.0 |
| | NREE (Dark) | 1.0 | 1.0 | 1.0 |

ANCOVA, analysis of covariance; TEE, total energy expenditure; REE, resting energy expenditure; NREE, non-resting energy expenditure. *Statistically significant difference between groups.
